# Connecting surveillance and population-level influenza incidence

**DOI:** 10.1101/427708

**Authors:** Robert C. Cope, Joshua V. Ross, Monique Chilver, Nigel P. Stocks, Lewis Mitchell

**Affiliations:** School of Mathematical Sciences, The University of Adelaide, Adelaide, SA 5005, Australia; Discipline of General Practice, The University of Adelaide, Adelaide, SA 5005, Australia; Stream Leader, Data to Decisions Cooperative Research Centre (D2D CRC), Adelaide, SA, Australia

## Abstract

There is substantial interest in estimating and forecasting influenza incidence. Surveillance of influenza is challenging as one needs to demarcate influenza from other respiratory viruses, and due to asymptomatic infections. To circumvent these challenges, surveillance data often targets influenza-like-illness, or uses context-specific normalisations such as test positivity or per-consultation rates. Specifically, influenza incidence itself is not reported. We propose a framework to estimate population-level influenza incidence, and its associated uncertainty, using surveillance data and hierarchical observation processes. This new framework, and forecasting and forecast assessment methods, are demonstrated for three Australian states over 2016 and 2017. The framework allows for comparison within and between seasons in which surveillance effort has varied. Implementing this framework would improve influenza surveillance and forecasting globally, and could be applied to other diseases for which surveillance is difficult.

## 1 Introduction

Influenza is a respiratory virus of substantial public health concern globally, causing variable morbidity and mortality depending on the strains circulating each season. Consequently, influenza is subject to surveillance in many jurisdictions. Influenza surveillance is difficult due to: the potential for mild or asymptomatic cases being unreported; influenza needing to be demarcated from other respiratory viruses in clinical settings; and, the complexity of influenza immunity (particularly relating to different strains and antigenic drift) [1, 2, 3, 4]. Due to this, influenza surveillance is often framed around influenza-like-illness (ILI) rather than influenza specifically, and reported under context-specific normalisations (e.g., per-consultation rates) that work in unintuitive ways and do not appropriately quantify uncertainty. This has unintended consequences, such as influenza estimates depending upon the prevalence of other respiratory infections. When influenza specific test results are reported, they often lack the denominators necessary to produce accurate quantitative comparisons within and between seasons [5]. Multiple surveillance data streams generally exist, but they are not assimilated to produce a population-level probabilistic estimate of influenza incidence. In the worst case the lack of such a framework could impact decisions based upon surveillance, such as the allocation of public health resources.

We propose that, when influenza is the primary concern, surveillance should strive to report, model, and assess influenza specifically, rather than ILI, at the population level. This framework can be achieved by carefully considering how population processes lead to observed data, i.e., the observation process, and how this introduces uncertainty to an analysis.

In addition to influenza surveillance, which addresses past and current influenza incidence, substantial research has been directed towards forecasting influenza incidence [6, 7, 8, 9, 10], including ongoing initiatives supported by the US CDC [11, 12]. Effective forecasting assists public health planning and management, providing estimates in advance for resource allocation. However, challenges in influenza surveillance extend directly to forecasting, but this has not been fully acknowledged to date. The uncertainty inherent in disease surveillance impacts not just the forecast, but propagates to assessments of forecast accuracy. Historical estimates of quantities such as the peak week of a season should be *probabilistic* estimates rather than *point* estimates, and consequently forecasts of these quantities should be assessed against probabilistic estimates. We demonstrate this new framework applied to influenza surveillance, forecasting and rigourous forecast assessment (Figure 1) using data from an existing influenza surveillance system in Australia (the Australian Sentinel Practices Research Network, ASPREN; [13]).

**Figure 1:**
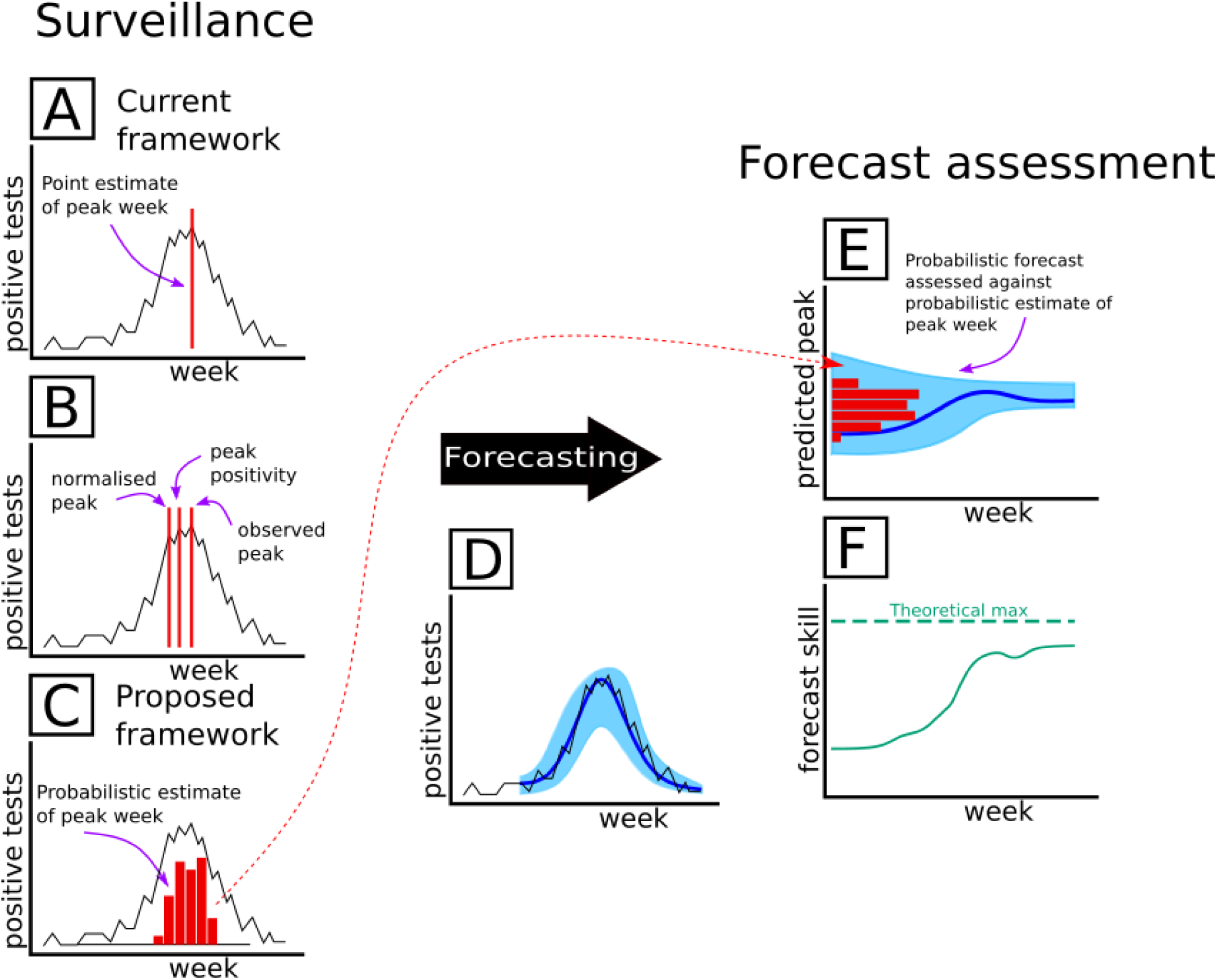
Schematic illustration of proposed framework for surveillance and forecasting, using peak week as an example. Given an observed influenza surveillance time series, the peak week is typically assumed to be a point estimate based on these data, as in (A); different ways of analysing these influenza surveillance data can lead to different point estimates of the peak week (B), for example based on test positivity, or per-consultation rates. Instead, we propose it should be a probabilistic estimate as in (C), because the observed time series is a sample or an estimate of the population-level dynamics, with associated uncertainty. Taking the observation process into account we can obtain probabilistic forecasts of influenza in the population (D), which can then be used to produce probabilistic forecasts of quantities such as the peak week (E). To assess these forecasts, we propose modified metrics that take into account that the true peak week has a distribution rather than a point estimate (F).

Our gold standard influenza surveillance and forecasting framework shifts to whole population estimates of influenza informed by observation processes, incorporating the variability inherent in surveillance systems; this also facilitates rigourous assessment of forecast outputs. This framework could also be applied more widely across communicable diseases, particularly when not every case is observed, tested or reported.

### 1.1 Influenza surveillance

Influenza surveillance systems are continually evolving, due to a combination of technological advances, new approaches, and renewed interest in response to the 2009 pandemic [14, 15]. Consequently, there is substantial variation in reporting at both long and short timescales (e.g., in outpatient or general practice based surveillance systems, such as the ILINet system in USA, or ASPREN in Australia [13]), or participation (in participatory surveillance systems, such as the Influenzanet program in Europe [16], or the Flutracking program in Australia [17, 18]). Further variation exists because some surveillance systems assess influenza specifically (requiring tests to be performed, possibly of variable accuracy, and where negative results may not be reported), while others measure influenza-like-illness (ILI) [5]. It is widely acknowledged that due to this variation (and in particular due to increases in surveillance effort over time), raw counts of ILI cases, or of positive tests, are not appropriate measures of population epidemic severity [5, 19]; to compensate, common strategies include: per-consultation or per-participant rates; test positivity; or, combinations of these strategies. These are intuitive and effective approaches; however, these highly-filtered observations and proxies provide an indication of the target quantity (i.e., population-level influenza incidence) that is subject to uncertainty. The relationship between observed data and population-level disease incidence should be described formally, and its uncertainty quantified. When the number of samples is very large, uncertainty may be relatively small, but should still be quantified, and often surveillance systems operate in contexts where only a small proportion of cases are tested, or only a small proportion of doctors report, leading to highly-filtered data and thus high uncertainty.

A further challenge with per-consultation or per-participant estimates is that they necessarily depend on additional factors that may have no relation to influenza and in addition may be poorly resolved: for example, participants in a voluntary online surveillance system may choose whether or not to report in a non-random way, or consultations at outpatient doctors may vary based on incidence of unrelated diseases (and so an outbreak of e.g. Gastroenteritis might inadvertently impact perceived influenza season severity). To rectify these problems, we propose that estimates from observed data should (carefully) drive probabilistic estimates at the population level, by considering, in each case, the (hierarchical) observation process that leads true population-level influenza incidence to be filtered down to the observed level. This careful analysis of the observation process should be applied at each stage of an analysis: during surveillance and reporting, and extended to modeling, forecasting, and forecast assessment.

### 1.2 Example surveillance system and observation process

The Australian Sentinel Practices Research Network (ASPREN) [13, 20] is a network of general practitioners (GPs) distributed throughout Australia. The target coverage is approximately one GP per 200,000 people in metropolitan areas, and one GP per 50,000 people in regional areas. This database is continually updated and requires the voluntary participation of GPs, so there is variation between years and locations. All GPs in the database report the number of consultations they performed each week, along with every case of a notifiable disease that they observe, including influenza-like-illness (ILI). GPs are asked to take swab samples from a proportion of those patients presenting ILI symptoms (20% in 2015-2017), and these samples are sent for virological (PCR) testing, resulting in respiratory virus virology data for each sample. In practice, not all GPs take samples, and the proportion of patients for which samples are taken varied substantially, however the number of doctors participating, and the number of ILI patients observed and tested is known.

The hierarchical observation process is adapted from Cope *et al*. [21], with key details re-stated here for clarity.

We proceed by considering how an individual in the population who is infected may come to be observed in the dataset. Filtering occurs at three levels:

- Not all individuals infected with influenza will seek treatment, as cases may vary in severity, including potentially being mild or asymptomatic. We assume that each infectious individual has some (unknown, independent and identically distributed) probability of seeking treatment.
- Only a small proportion of GPs are part of the ASPREN network, and this proportion has varied over time. We assume that if an infectious patient consults a GP, then that GP has some fixed probability of being part of the ASPREN network (estimated annually based on the number of doctors that participate).
- The proportion of ILI cases that are sent for testing varies substantially (see Figures S1-3), both due to changes in ASPREN protocols, and variation in doctor behaviour. We assume that each infectious patient who attends an ASPREN GP has some probability (fixed, estimated from data, varying weekly) of being sent for testing.

To obtain population-level estimates from these data, we model this observation process: if *Z_j_* is the number of individuals in the population seeking treatment for symptomatic influenza in week *j (j ∈* {1, 2,…, 52}), then we observe *n_j_* ~ Bin(*Z_j_, p_j_*) confirmed cases, with *p_j_* the normalization probability (i.e., the proportion of doctors in the state that are in the dataset, multiplied by the proportion of ILI cases those doctors sent for testing in that week). So, given *n_j_* and *p_j_*, the number of symptomatic individuals is *Z_j_* ~ NegBin(*n_j_* + 1, *p_j_*). Critically, this is a probabilistic estimate rather than a point estimate (e.g., Figure 2). Note that this is an estimate of the number of individuals who seek treatment with influenza; to obtain the total number of individuals with influenza we would also require an estimate of the probability of seeking treatment, which is not directly available from these data (though can be estimated using a model).

**Figure 2:**
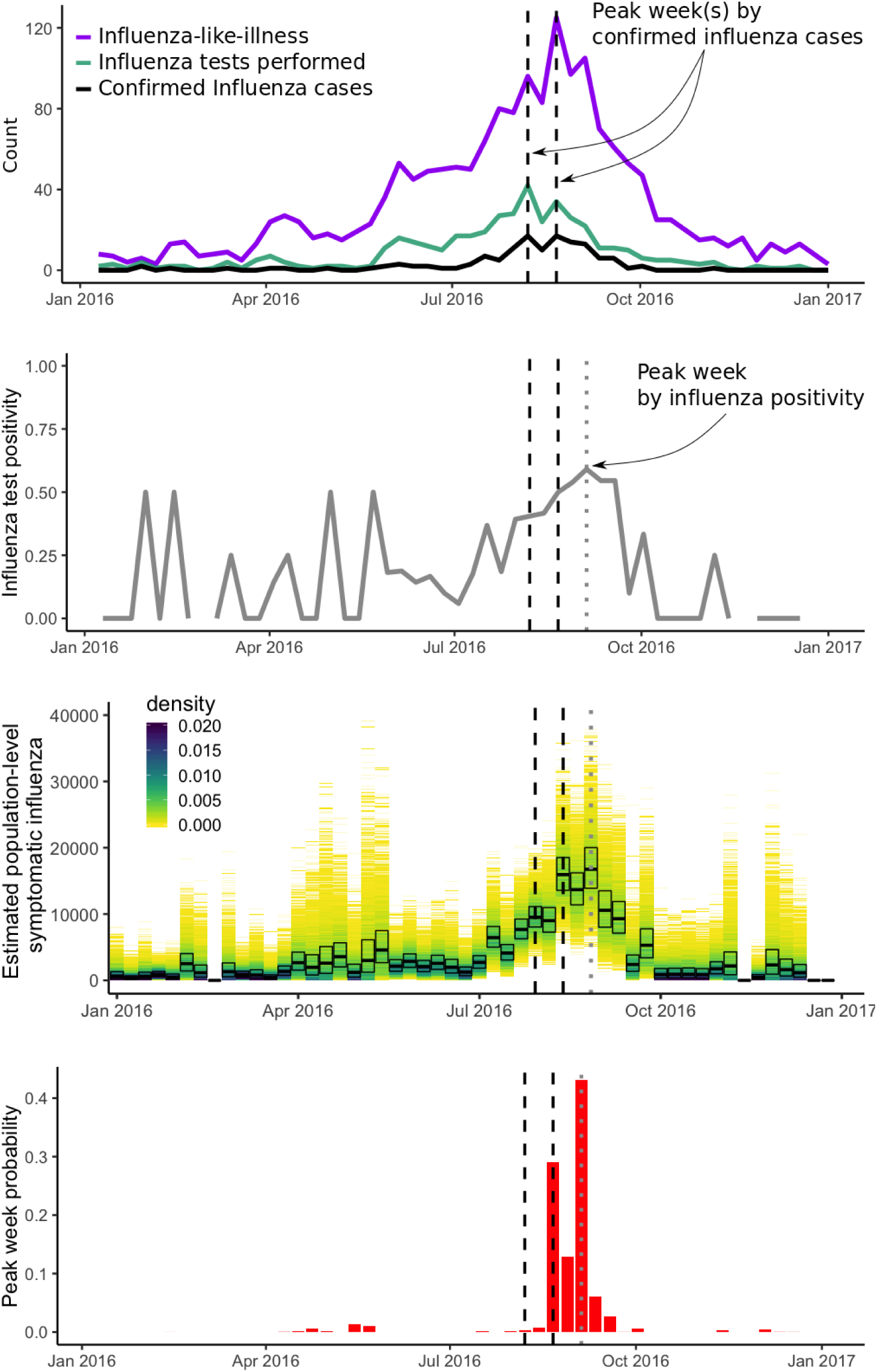
2016 ASPREN Influenza surveillance data from New South Wales, demonstrating how observed data-level peaks can be misleading. The top figure shows confirmed influenza (black) and ILI (purple), along with the number of samples tested each week (green). These data would suggest peak-week point estimates in the weeks ending 6 August and 20 August (dashed black lines). The second figure (grey) shows influenza test positivity, which peaks during the week ending 4 September (dotted grey line), and is highly variable outside of the season when few tests were performed. The third panel (yellow - blue colour gradient) shows estimates of population-level symptomatic influenza incidence each week, with uncertainty: black boxes indicate the median and a 50% credible interval, with the shading showing the full density. The lowest panel (red) shows the distribution of population-level peak week calculated using all data. This demonstrates that the peak week inferred directly from the surveillance-level data can be misleading, and that test positivity is also a flawed metric.

### 1.3 Influenza forecasting

In contrast to surveillance uncertainty, practitioners understand that forecast uncertainty is critical, and best practice includes reporting and assessing influenza forecast uncertainty ([12, 22]). A consequence of overlooking uncertainty in surveillance data is that, particularly when data are highly filtered or stratified (resulting in substantial uncertainty), the absence of uncertainty in estimates flows through to an absence of uncertainty in subsequent analyses. For example, assessing forecasts against point-estimate surveillance data could result in erroneous assessments; rather, forecasts should be assessed against probabilistic estimates of the true epidemic behaviour. Incorrect forecast assessment could lead to choosing inferior forecasting methods to inform management decisions, which could in the worst case have detrimental public health impacts, for example, planning hospital bed availability to peak at the wrong time.

A quantity of interest for surveillance and forecasting is the peak week, i.e., the week during the season that influenza incidence is highest. When the population-level influenza estimate for each week has a probability distribution, then the peak week must also have a distribution (i.e., the distribution of the maximum of the *Z*_1_*,…, Z*_52_). This can be evaluated through simulation. Figure 2 shows the relationship between surveillance data, test positivity, and population-level peak week distribution for data from New South Wales in 2016, demonstrating that considering the data directly or relying on positivity alone can be erroneous and at least fails to quantify the uncertainty inherent in estimates of peak week from surveillance data. Producing population-level estimates of symptomatic influenza incidence (Figure 2c) combines the ILI data, the number of tests performed, and the results of those tests to quantify incidence and uncertainty through the season; which is then used to produce a probabilistic estimate of the peak week (Figure 2d). After estimating this distribution for the true peak week, we can use it to assess forecasts, by generalizing metrics such as forecast skill, to average forecast skill over the distribution (see Section 4.6 for methods).

## 2 Results

To illustrate the link between influenza surveillance, forecasting, and forecast assessment, we apply this framework with ASPREN surveillance data in three Australian states (New South Wales, Queensland, and South Australia) for 2016 and 2017. We use surveillance data and the observation process to evaluate population-level symptomatic influenza incidence estimates, and use this to evaluate a distribution for the peak week (Figures 2, S1-S3). We used a two-stage approximate Bayesian computation based forecasting framework (see Section 4.3.3), with an underlying population-level stochastic epidemic model and an observation process explicitly linking surveillance data to population-level influenza (e.g. Figure 3). We extracted forecast distributions for peak week, which are assessed against probabilistic estimates of true peak week at the population level (Figure 4). We observed that forecasting was possible when data were sufficient, such as in New South Wales (both years), and in 2017 for Queensland and South Australia; but when data were very sparse it was not possible to effectively produce or assess forecasts (e.g. for South Australia in 2016). We evaluated average forecast skill (see Section 4.3.4) each week, and observed higher forecast skill in 2017, with the best performance in New South Wales (Figure 5). Due to the sparsity of data, the best achievable forecast skill was lower in 2016 than 2017, and particularly low in South Australia.

**Figure 3:**
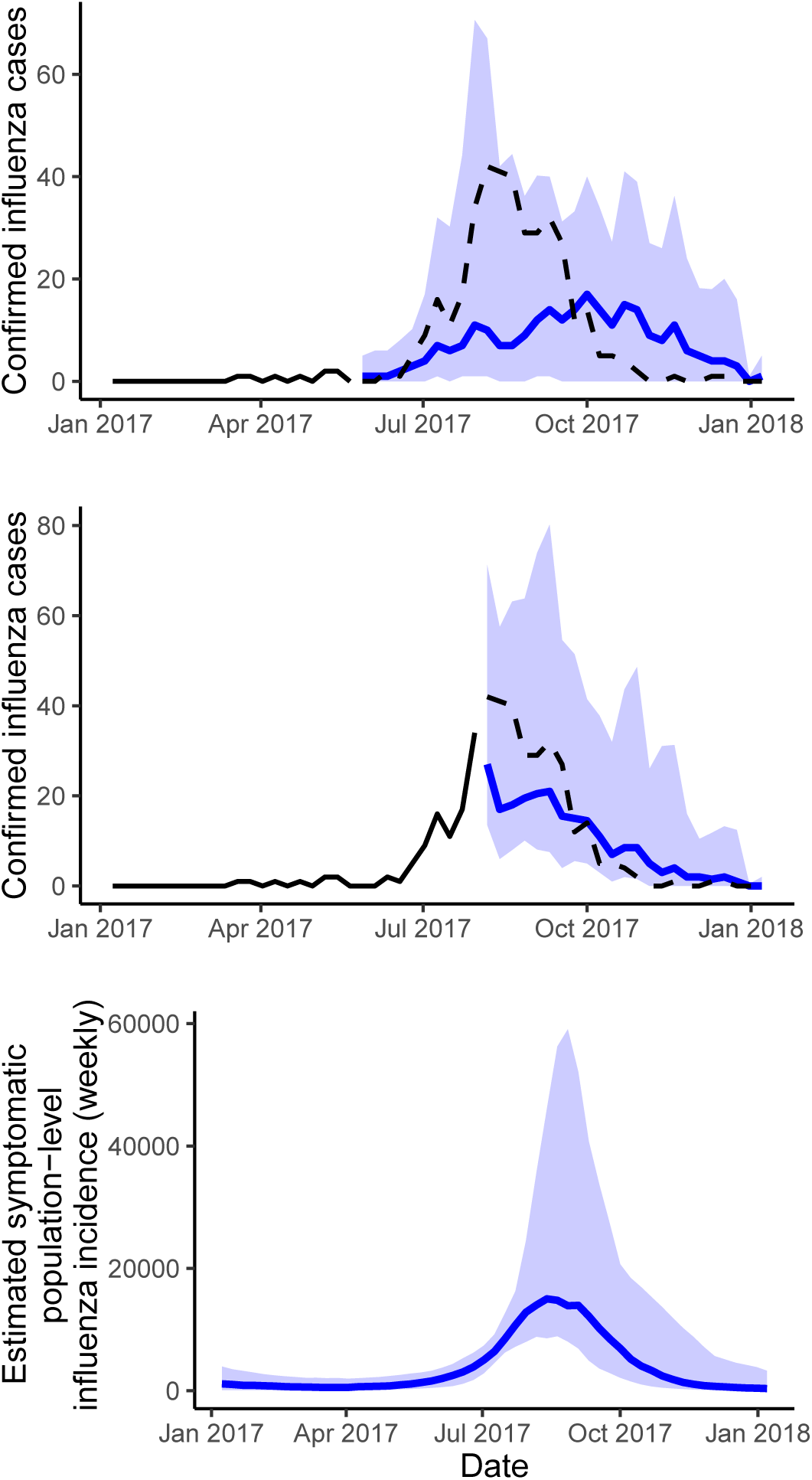
Forecasts for New South Wales in 2017: (top) before the start of the season, projected down to data-level, (middle) during the season, projected down to data-level, and (bottom) during the season, at population level. The black line is the observed data; the blue line the median forecast, and the shaded area the 90% prediction interval. Example forecasts projected down to data-level for all states and all years appear in the Supplemental Information (Figures S4-S5).

**Figure 4:**
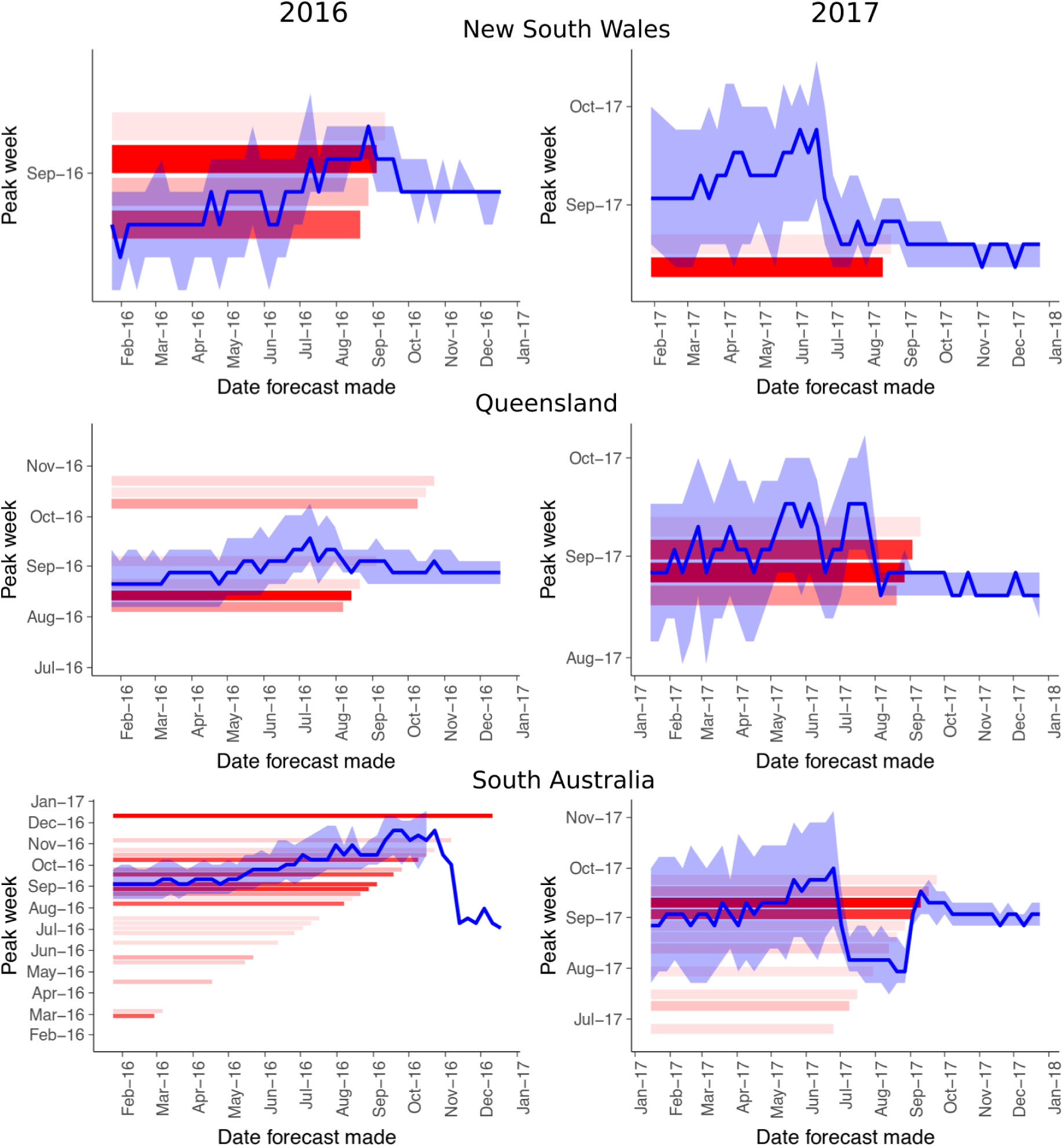
Progression of peak week forecasts each week during the season for New South Wales (top), Queensland (middle), and South Australia (bottom) during 2016 (left) and 2017 (right). The blue line indicates the median forecast, and the light blue area the 50% prediction interval. Red bars indicate the distribution of the population-level peak week, with the alpha proportional to the probability mass (i.e., darker bars correspond to higher probability); only those weeks with probability ≥ 0.05 are shown. The length of the red bars indicates where the axes are equal, i.e., the forecast is the current week. Therefore, predictive forecasts should converge to a given value before the end of the bar. See Appendix for influenza time series (Figures S1-S3) and example forecasts (Figures S4-S5).

**Figure 5:**
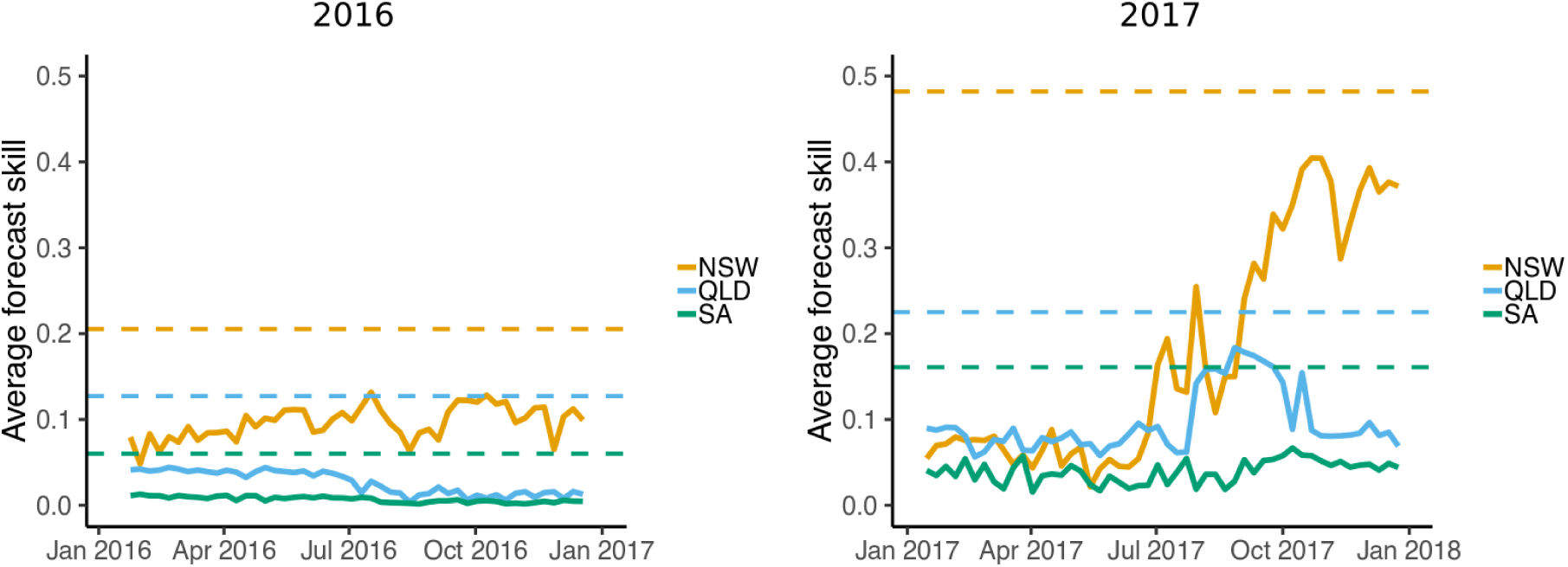
Progression of average forecast skill each week during 2016 and 2017. Note that while forecast skill can theoretically reach 1.0, this requires ideal conditions; in practice, the best achievable forecast skill would be that achieved by the observed distribution of peak week from the data shown here as a dotted line. See Appendix for influenza time series (Figures S1-S3).

## 3 Discussion

Influenza surveillance and forecasting can provide important information for public health decision making. Of primary interest is the incidence of influenza, specifically, within populations. Unfortunately, current approaches to surveillance and forecasting do not evaluate this. Instead, they often work with influenza-like-illness (ILI), and operate at variable scales influenced by a myriad of other factors. In this study we demonstrate that analysis can be performed on influenza specifically, at the population level, by carefully considering the observation process and denominators that link population-level processes to observed data. Estimating influenza incidence at the population level allows comparison between historical seasons during which different numbers of tests were performed, and reduces the impact that incidences of other viruses have on influenza surveillance. As a specific example, Lambert *et al*. [5] describe how a hypothetical outbreak of respiratory syncytial virus could bias influenza estimates based on test positivity: under our framework, this potential bias would be eliminated. This shift in framework not only impacts surveillance and forecasting, but also allows for rigourous assessment of forecast accuracy that takes into account uncertainty in estimates of true data.

While we claim that, in most cases, the goal in influenza surveillance and forecasting should be to analyse population-level influenza incidence itself, it is possible that in some cases there may be other targets of interest. For example, in some applications one may be interested in influenza-like-illness presentations generally at particular points of care, such as at emergency departments. In this case, it would be reasonable to surveil, report, and forecast that quantity of interest, specifically. We advocate for careful consideration of the quantity of interest, how that links to the data available, and direct evaluation of that target quantity and its uncertainty.

A challenge posed by this shift in framework is communicating these ideas to the public health community. We claim that operating at the population level and with confirmed influenza specifically is an intuitive goal, but acknowledge that changes in approach, and in particular introducing uncertainty around perceived “truth” values such as the peak week, will require a concerted effort led by uptake by informed practitioners. But this shift in framework to surveilling and forecasting the quantities of interest will be of substantial benefit to the public health community.

### 3.1 Probabilistic forecast assessment

Recognizing that perceived “truth” values are themselves estimates with associated probability distributions, rather than point values, we require new, rigourous metrics for assessing forecast accuracy. There are a range of possible metrics; we chose examples that are natural generalisations of metrics currently used for assessing forecasts around point estimates [11, 12], by averaging over the distribution of the true peak week. However, two challenges remain: (i) since this approach to forecast assessment is new, there are no existing results to compare to, and (ii) metrics that assess forecasts against probability distributions depend on the probability distribution for the “truth”, such that no forecast can ever reach a “perfect” score (e.g., a forecast skill of 1, or average error of 0; except when the “truth” is a point estimate). There are substantial differences between the best possible scores in different states or years (Figure 5). For this reason, we suggest that metrics such as averaged forecast skill should be used as tools for comparison between methods (or over time), when comparing in the same location, against the same estimated truth distribution.

One tangential advantage of these metrics is that it allows one to retrospectively assess the value of the data being used for forecasting, by comparing the scores that would be achieved by the truth distribution itself. We observe this in the current study (Figure 5), whereby maximum possible forecast skill for peak week is substantially higher in 2017 than 2016 in all locations due to more data and clearer peaks. Maximum possible forecast skill for peak week was lowest in South Australia in 2016 where data was insufficient to perform effective forecasting, and there was no clear peak in the data.

We do not expect the distribution of a forecast *(p_w_*, the forecast probability that the peak is week *w*) to resemble the distribution of the truth (*q_w_*, the probability that the peak is week *w* given the observed data), given that they are fundamentally different quantities. The distribution of the forecast takes into account all of the data from the season to produce a posterior forecast of epidemic trajectories (which becomes tighter as more data becomes available). Whereas the distribution of the truth treats each week as an independent sample of the unobserved population-level quantity.

Note that point estimates of disease incidence can artificially inflate forecast accuracy score, because once, say, the peak week is observed, its value is known exactly. As such, if you are certain that you have observed the peak, your forecast can assign all probability to that week. This is why, for example, the forecasts assessed in Biggerstaff *et al*. [12] rapidly achieve skill scores of 1.0 after the peak has occurred.

### 3.2 Assumptions, limitations, and future work

One challenge of the forecasting we use is that to produce forecasts of influenza requires forecasts of temperature. In this study, we used a simple statistical time-series approach; it may be the case that improved forecasting of temperature could be obtained with more complex methods or by using externally-produced forecasts (i.e., from the Bureau of Meteorology; www.bom.gov.au). However, it is likely that stochasticity in both the disease and observation processes outweighs variation in temperature forecasts.

One limitation of this work is that it considers only a single influenza surveillance data stream. In practice there are many complementary surveillance systems, and while none assess population-level influenza incidence directly, they each provide useful information. Future work will develop a rigourous framework to assimilate a full range of disease surveillance data streams to produce a combined estimate of population-level influenza incidence.

### 3.3 Conclusion

Disease surveillance, and subsequent analyses such as forecasting, should be focused on the quantity of interest, at the scale of interest; for influenza, this means evaluating influenza incidence itself, at the population level. We demonstrate a framework in which this is possible, and show that this impacts not just forecasting, but also for the assessment of forecasts. This framework should be applied broadly by researchers and public health professionals, both for influenza and more generally across communicable diseases.

## 4 Methods

### 4.1 Data

Influenza data were obtained as part of the ASPREN project (13; https://aspren.dmac.adelaide.edu.au/). In this study, we used those data from three states: New South Wales, Queensland, and South Australia. We produced forecasts for the 2016 and 2017 seasons, using the two preceding years of data to inform prior distributions in each case. For this to be possible, we required sufficient levels of epidemic data from the two preceding years, as sparse historical data could produce poor estimates. Forecasts were based on weekly aggregate counts of influenza, tests, ILI, and the number of doctors contributing to the database.

Historical climate data were obtained from the Australian Government Bureau of Meteorology (www.bom.gov.au/climate/data), at stations 066062 (Sydney - Observatory Hill), 040913 (Brisbane), and 023090 (Adelaide - Kent Town). We used climate data from state capital cities in each case, as both the majority of the state population and the majority of doctors in the study were located in these cities. Temperature observations were taken every three hours (i.e., 8 times per day). Note that three-hourly observations creates a cyclic temperature pattern with both daily and annual periods. So as to avoid having unrealistic diurnal dynamics, we used the average daily temperature rather than the three-hourly observations themselves.

We used fixed, total population sizes of 6.9 million, 4.3 million, and 1.6 million, in each state respectively, based on 2011 Australian Bureau of Statistics estimates. While in practice these populations are increasing over time, including demography would add considerable complexity, which we wished to avoid.

### 4.2 Estimating population-level peak week from surveillance data

Using surveillance data and the observation process, we can evaluate the distribution of population-level influenza incidence, each week. We can compare these distributions across a year to produce a distributional estimate of the peak week. This proceeds as follows:

- For each week, given *n_j_* observed influenza cases with *p_j_* the normalization probability (i.e., the proportion of doctors in the state that are in the dataset, multiplied by the proportion of ILI cases those doctors sent for testing in that week), generate *Z_j_* ~ NegBin(*n_j_* + 1, *p_j_*).
- Compare the generated *Z_j_* to determine the peak week.
- Repeat this process many (e.g., 10,000) times and collate the results to evaluate the distribution for the population-level peak week.

### 4.3 Influenza forecasting

#### 4.3.1 Epidemic model

We constructed a stochastic epidemic model for the spread of seasonal influenza within a population (Figure S6; following [21]). This model featured temperature-driven transmission, with 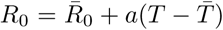, where T is the mean daily temperature, 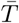 is the mean over the full temperature dataset, and Ro and *a* are fitted parameters (using approximate Bayesian computation; see Section 4.3.3, and Supplementary Materials). The model operated continuously through time, across years with waning immunity, rather than modelling each season separately, and treating influenza as a single disease (rather than separating strains). This means that consecutive seasons are related, and that at the beginning of a season there remains immunity in the population due to transmission in previous years. Rather than evaluate exact simulations of a full continuous-time model, a binomial approximation was used with 8 timesteps per day (i.e., corresponding to the climate observations).

#### 4.3.2 Temperature forecasting

To forecast influenza using a model that is driven by temperature, we were first required to forecast temperature dynamics, at a three-hourly resolution (i.e., 8 observations per day). Our approach was to use statistical time-series methods, in particular an autoregressive integrated moving average (ARIMA) model with seasonal forcing at both annual and daily timescales. We used the ‘forecast’ package in R [25, 26] to fit Fourier *x*—regularisation terms to capture seasonality, and to fit an ARIMA(1,0,1) model to temperature, from which we could then simulate. This produced a seasonal pattern that appeared similar to historical temperature data, although had fewer extremely hot or cold days. While this model is not perfect, it appeared to sufficiently capture the seasonality in temperature which would then drive the seasonality in influenza dynamics.

#### 4.3.3 Forecasting algorithm

Assume that we wish to produce a forecast from data up until week *w* of year *Y* (Figure 6). For all preceding weeks, assume we have full climate information, weekly influenza data, and denominators (i.e., the number of doctors participating, and the proportion of ILI cases sent for testing). Influenza forecasting proceeds as such:

1. Fit the model to all of the observed data from years *Y* − 2 and *Y* − 1, using approximate Bayesian computation and an uninformative prior (ABC; see Supplementary Materials for details). The result of this process is a posterior distribution, in the form of a collection of particles, which includes both the model parameters, and the (unobserved) model states. This posterior distribution is retained as a collection of particles.
2. Consider the data from the current year, for weeks 1,…, *w*. Use ABC to fit the model to these data. The prior distribution for this process is a modified version of the final posterior distribution from Step (1), including initialising the unobserved model states at the beginning of this year as those from the posterior distribution at the end of the previous year.

- To modify the prior so as to allow for between-season variation due to, e.g., differences in strains: (i) select a random time point, uniformly, from week 20 to week 27; (ii) at that time point, reallocate individuals between the (hidden) susceptible and recovered classes, by way of a mixture distribution, i.e., generate *u* ~ *U*[0,1]. Then the updated *S* and *R* values (denoted *S*\* and *R*)* are:

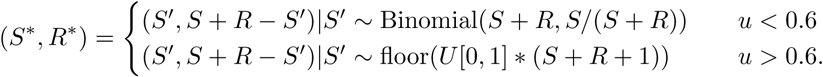 The mixture proportion (0.6) was chosen to balance the potential for both similar strains and shifts to substantially different strains.
3. Produce temperature forecasts for the remainder of the current season using an ARIMA model as described above.
4. Sample from the accepted particles and project forward from their state at the end of the ABC period. Retain the sample paths.

**Figure 6:**
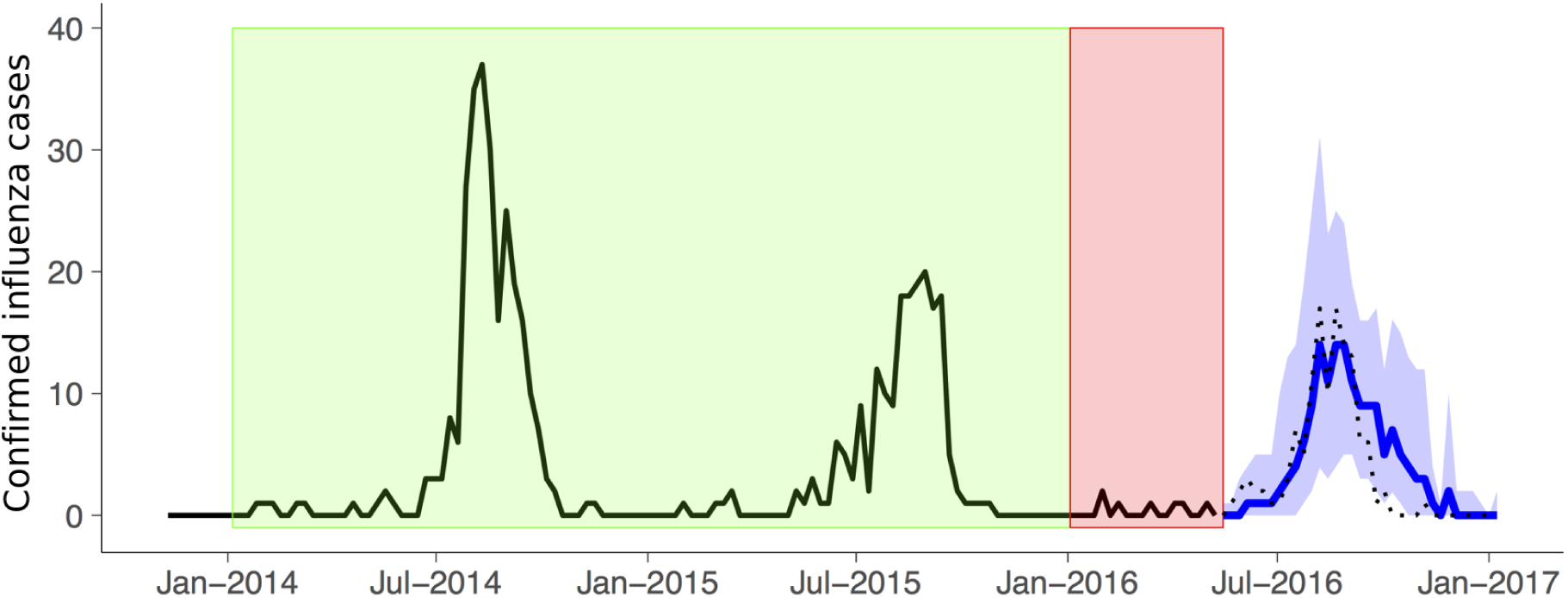
Schematic describing forecasting process for an example week in 2016. The black line is the observed data (dotted over the forecast period). ABC is performed on two preceding years of data (green region), and the posterior used as a prior for the current season. ABC is then performed again within the current season, based on data observed to date (red), and a forecast is produced for the remainder of the season by progressing accepted particles forward through the rest of the season (blue solid line is the median forecast; shaded region is the 50% prediction interval). The population-level forecast is projected down to the data-level using the true normalization data observed in that week.

### 4.4 Forecast assessment

Advances in epidemiological forecasting have been concomitant with advances in the reporting and assessment of forecasts. Historical forecast assessment most commonly assessed point forecasts directly, by asking if the point forecast was within some interval (e.g., ±1 week) of the observation (e.g. [11]). As emphasis has moved to probabilistic forecasting, forecast assessment has moved towards logarithmic scoring rules (e.g., [12]). Held and Meyer [22] provide a comprehensive summary of the state of the art in forecast assessment.

Let *p_w_* be the forecast probability that the peak week is week *w*, and *q_w_* be the probability that *w* is the true population-level peak week. We report two accuracy metrics:

- Forecast skill score, i.e., the exponential of log score average over the true week distribution: 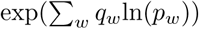. This is equivalent to the forecast skill score used elsewhere [12] when the truth is a point estimate (i.e., only has density in a single week).
- Average error in peak week predictions: 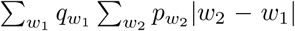. This is equivalent to average (absolute) error if the truth is a point estimate. This is reported in the Supplemental Information (Figure S7).

There are other metrics that could reasonably be used (e.g., mean squared error, or Kullback-Leibler divergence), however we chose to focus on these two as they are most analogous to those used in existing forecasting analyses.

## Acknowledgements

**General:** We thank the general practitioners and nurse practitioners whom report data to ASPREN, SA Pathology for laboratory testing, Adelaide Health Technology Assessment for database management and Datavation for FluSync tool development.

## Author contributions

R.C.C., J.V.R. and L.M. designed the project. M.C. and N.P.S. collected and curated the influenza surveillance data. R.C.C. performed the analysis and drafted the manuscript. All authors edited the manuscript.

## Data availability

Upon publication, data will be made available at a public repository.

## Supplementary Materials

### Example data streams

**Figure S1:**
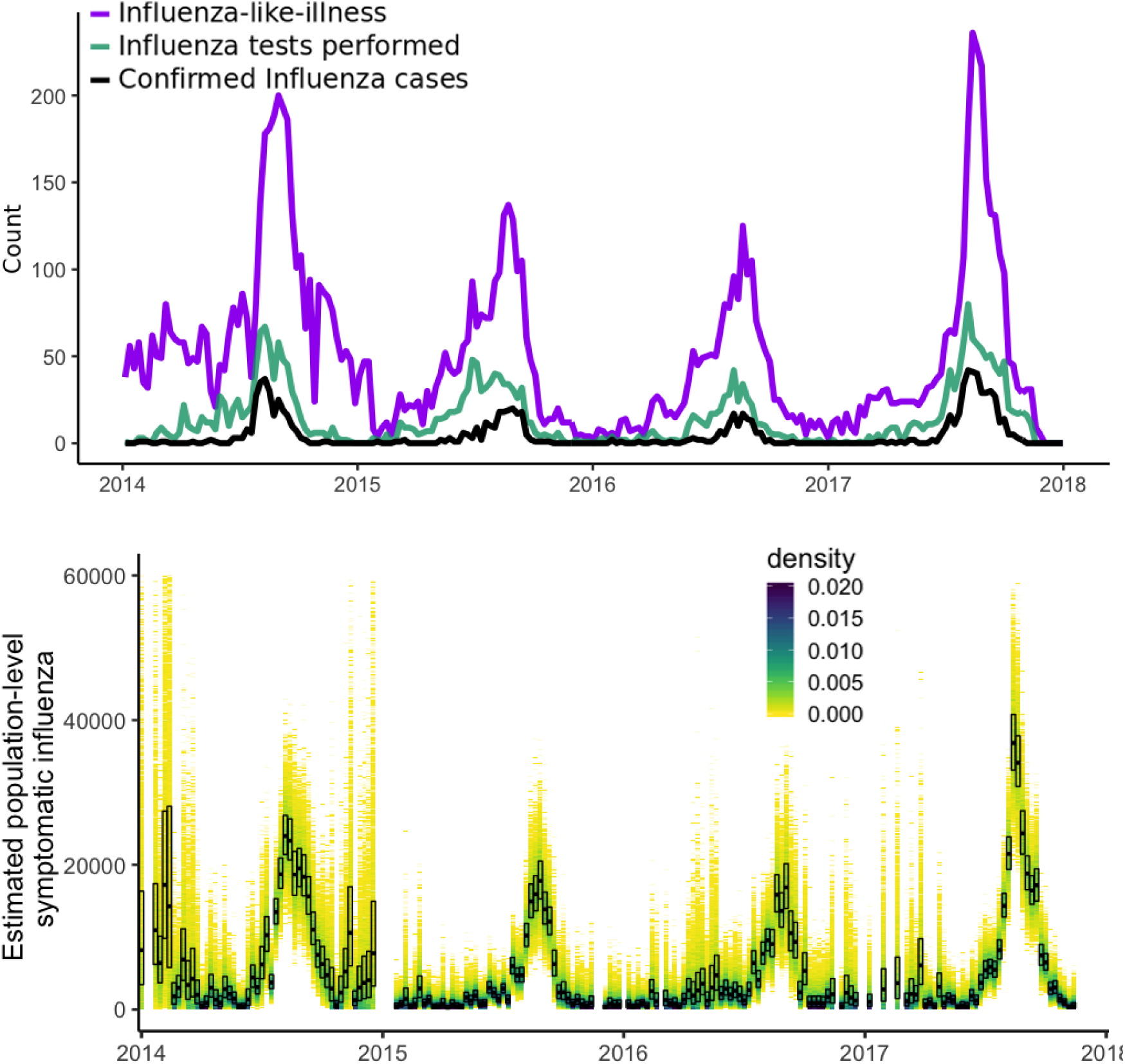
(above) Confirmed influenza (black), testing (green) and influenza-like-illness (purple) in New South Wales across 2014-2017, in the ASPREN database. Data from 2014-2015 were used to construct priors that informed forecasting in 2016; similarly, data from 2015-2016 were used to construct priors that informed forecasting in 2017. (below) Estimated population-level symptomatic influenza incidence in New South Wales across 2014-2017, based on the ASPREN data. Weeks in which no influenza tests were performed are omitted. Shading (orange) indicates probability density, boxes (black) indicate median and 25-75% quantiles.

**Figure S2:**
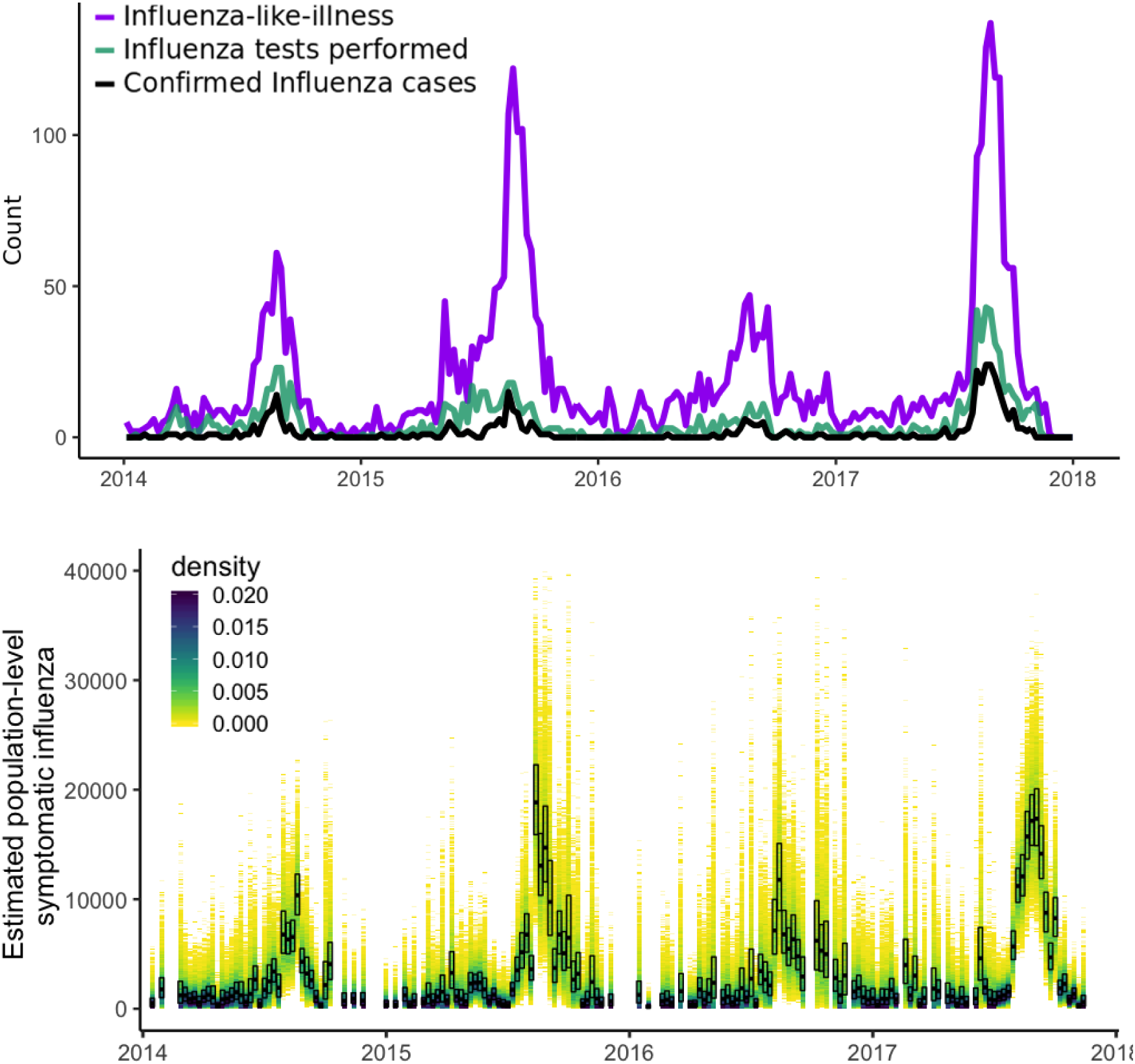
(above) Confirmed influenza (black), testing (green) and influenza-like-illness (purple) in Queensland across 2014-2017, in the ASPREN database. Data from 2014-2015 were used to construct priors that informed forecasting in 2016; similarly, data from 2015-2016 were used to construct priors that informed forecasting in 2017. (below) Estimated population-level symptomatic influenza incidence in Queensland across 2014-2017, based on the ASPREN data. Weeks in which no influenza tests were performed are omitted. Shading (orange) indicates probability density, boxes (black) indicate median and 25-75% quantiles.

**Figure S3:**
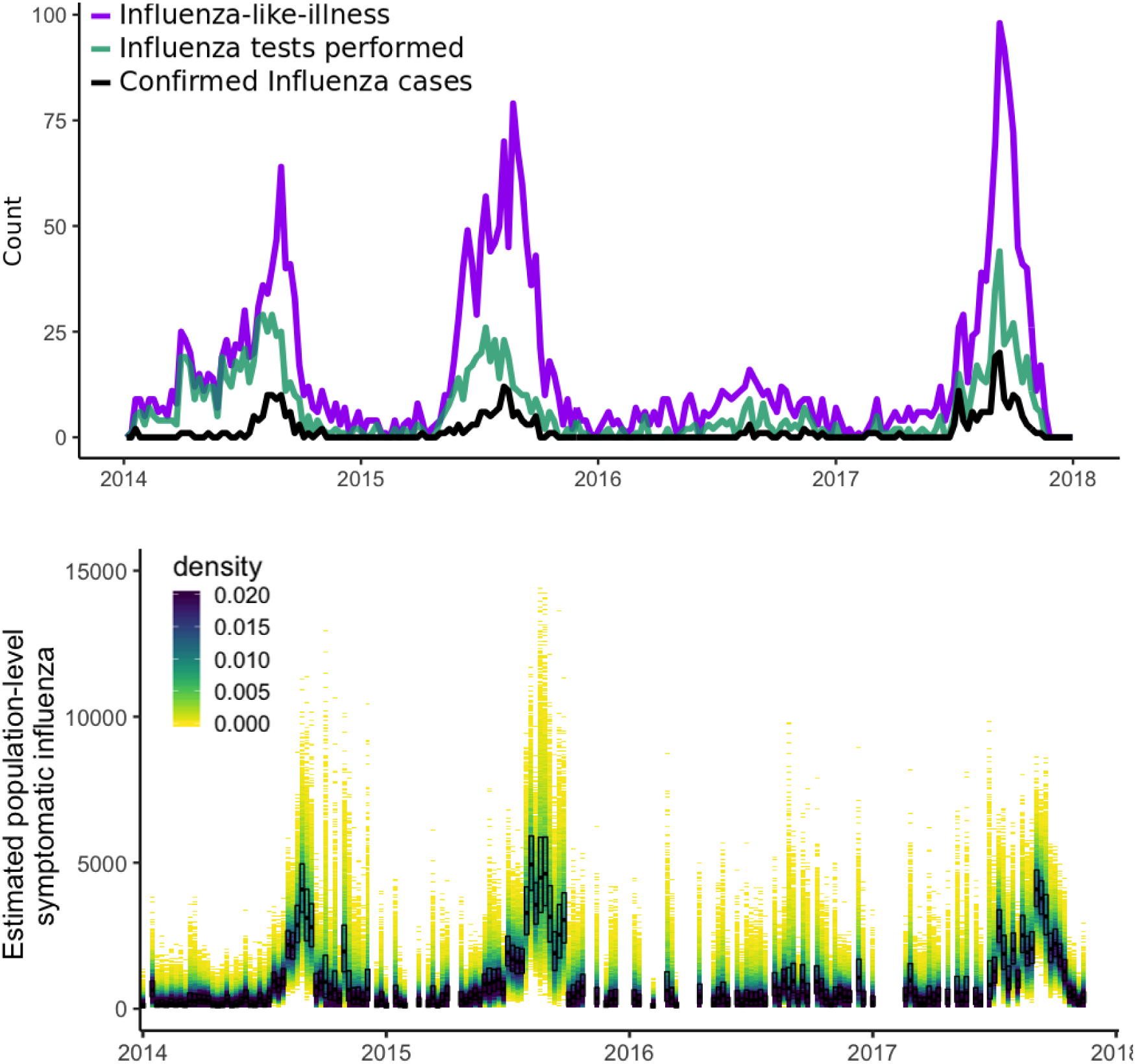
(above) Confirmed influenza (black), testing (green) and influenza-like-illness (purple) in South Australia across 2014-2017, in the ASPREN database. Data from 2014-2015 were used to construct priors that informed forecasting in 2016; similarly, data from 2015-2016 were used to construct priors that informed forecasting in 2017. (below) Estimated population-level symptomatic influenza incidence in South Australia across 2014-2017, based on the ASPREN data. Weeks in which no influenza tests were performed are omitted. Shading (orange) indicates probability density, boxes (black) indicate median and 25-75% quantiles.

### Forecast examples

**Figure S4:**
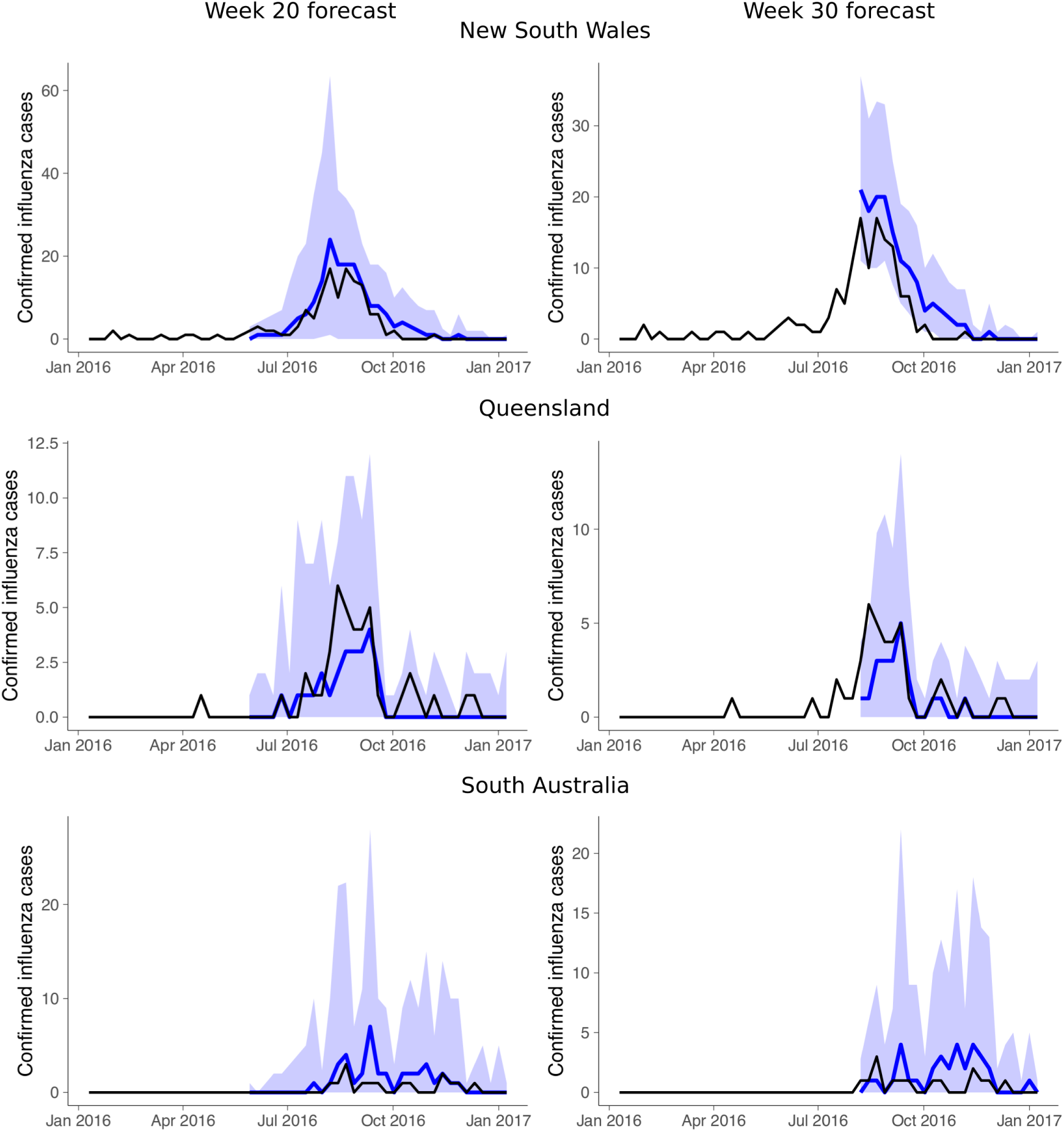
Illustrative forecasts from New South Wales (top), Queensland (middle), and South Australia (bottom) for 2016, projected down to the level of the data. The left panel shows forecasts made before the start of the season (week 20); the right panel forecasts made during the season (week 30). The red line shows the observed influenza data, the blue line the median forecast and the light blue area the 50% prediction interval.

**Figure S5:**
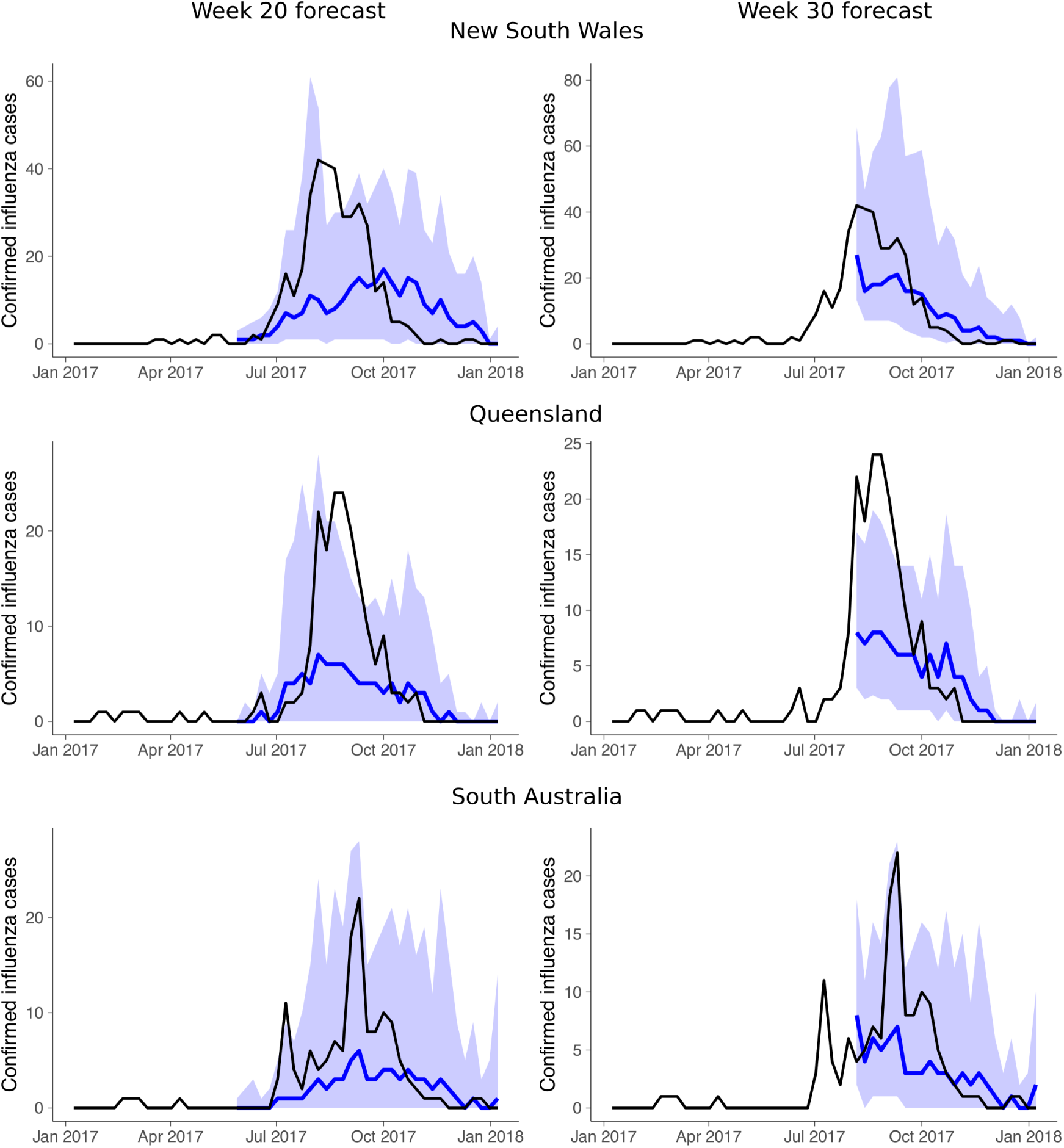
Illustrative forecasts from New South Wales (top), Queensland (middle), and South Australia (bottom) for 2017, projected down to the level of the data. The left panel shows forecasts made before the start of the season (week 20); the right panel forecasts made during the season (week 30). The red line shows the observed influenza data, the blue line the median forecast and the light blue area the 50% prediction interval.

### Approximate Bayesian computation (ABC)

Approximate Bayesian computation (ABC) [23, 24] is a likelihood-free Bayesian statistical method, which we use for parameter estimation as part of forecasting. We aim to estimate the (joint) posterior distribution for model parameters (e.g., rates of infection, recovery, waning immunity, treatment seeking) in the stochastic epidemic model (Figure S6; following Cope *et al*. [21]); to do this we select parameters sets (particles) from the prior, simulate from the model using the particle, and accept the particle as part of the posterior if the resulting simulated realisation (i.e., a time-series of weekly confirmed influenza counts at ASPREN doctors) is close to the observed data under a chosen metric. In this study, we chose a weighted squared error metric:

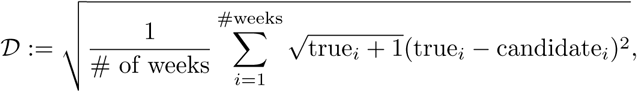

with true*_i_* the observed number of cases in the *i^th^* week from the ASPREN dataset and candidate*_i_* the number in the simulated realisation from the candidate particle. The weighting was chosen so as to value weeks with higher influenza activity (within the season) higher than those with little activity. When fitting the two preceding seasons to the season of interest, the threshold was set to 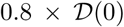, where 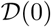 is the score that would be achieved by a season with no influenza activity. This was to ensure that the accepted realisations did actually resemble the data, while allowing for variation in activity between seasons. When fitting during the season, the threshold was chosen adaptively to ensure an acceptance proportion of 1%.

Prior distributions for the initial, two-year fitting period were chosen to be uninformative. Exponentially-distributed priors were used for latent, infectious, and immune periods, so as to have unbounded positive support (with means of 1 day, 2 days, and 10,000 days, respectively). Transmission was temperature dependant, with 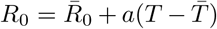; so 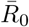 had a Gamma(2,2.5) distributed prior, and the seasonality parameter *a* was -Exp(0.2). The probability an infected individual would seek treatment had a Uniform(0.1,1.0) prior. In the second ABC phase, fitting to the season of interest, the prior distribution was taken to be the posterior sample from the initial two-year fitting.

Given that the model treats influenza as a single disease rather than fitting strains separately, prior and posterior distributions should not be treated as biologically informative, rather the goal of this analysis was solely to obtain forecasts. An epidemiologically-motivated analysis of influenza based on ASPREN data appears in Cope *et al*. [21].

**Figure S6:**
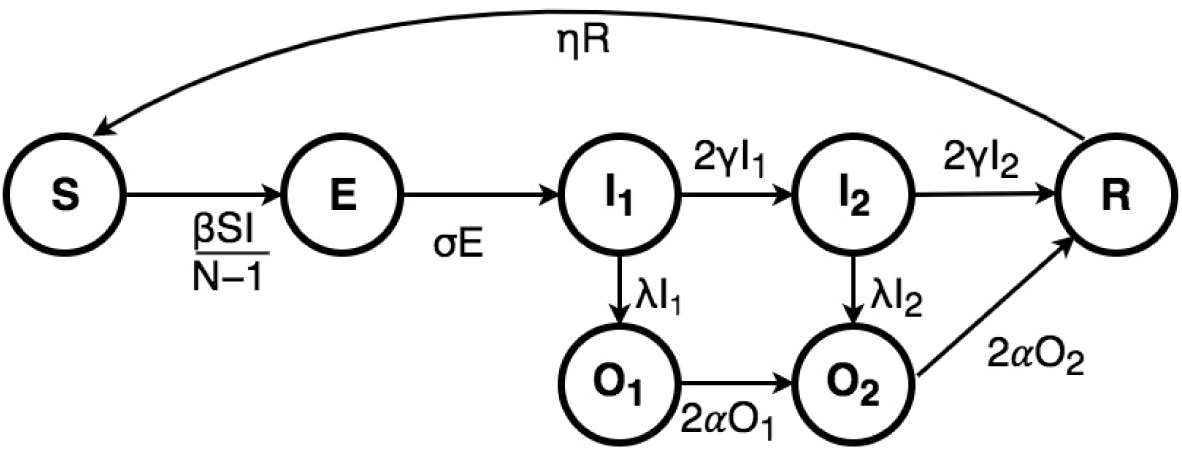
Stochastic epidemic model schematic.

### Average Forecast Error

**Figure S7:**
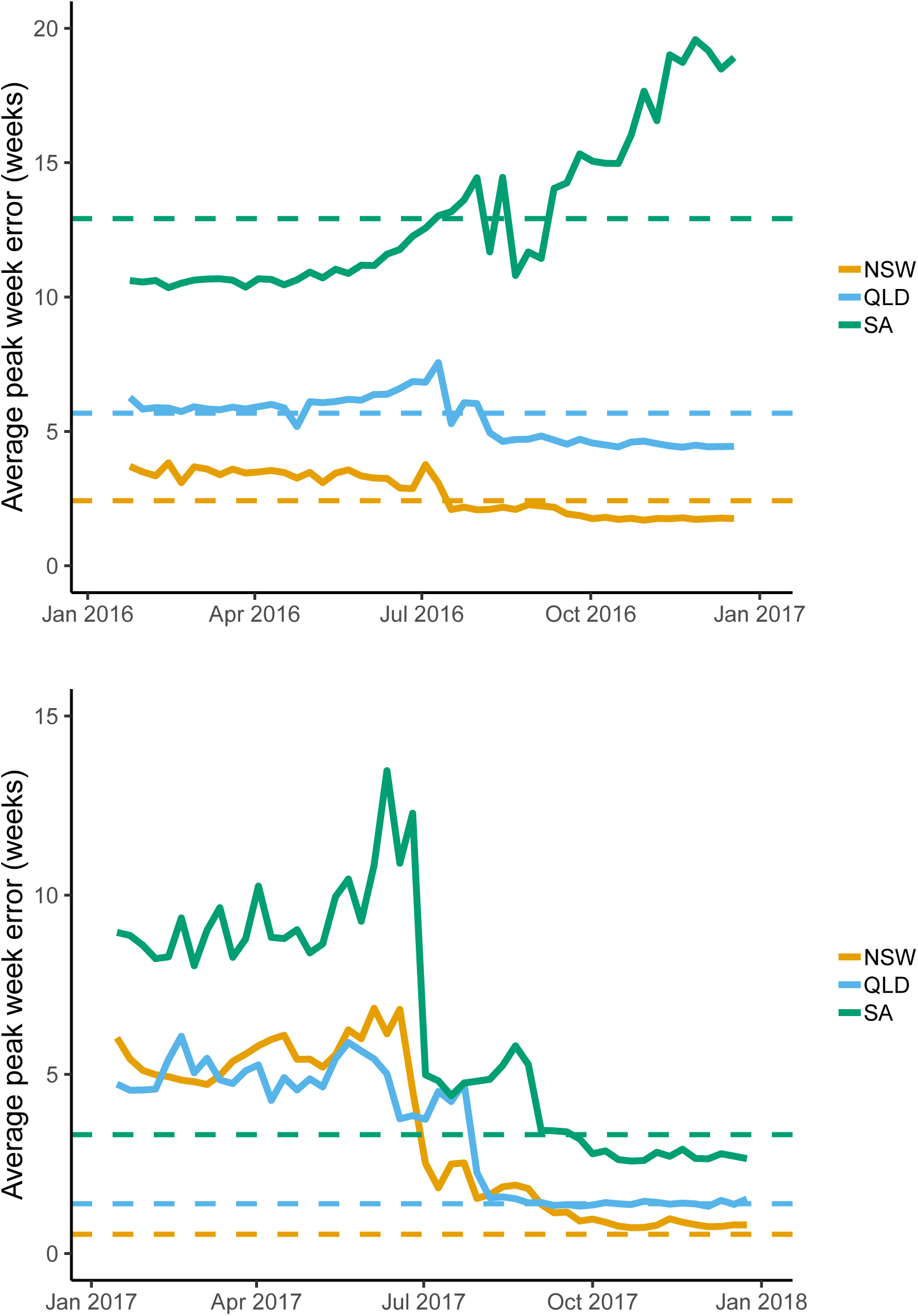
Progression of average forecast error each week during 2016 and 2017. Note that while forecast error can theoretically reach 0, this requires ideal conditions; in practice. An illustrative baseline of achievable forecast error is that achieved by the observed distribution of peak week from the data, shown here as a dashed line.

